# Association the 25(OH) vitamin D status with Upper Respiratory Tract Infections Morbidity in Water Sports Elite Athletes

**DOI:** 10.1101/559278

**Authors:** Jamshid Umarov, Fikrat Kerimov, Abdurakhim Toychiev, Nikolay Davis, Svetlana Osipova

**Affiliations:** Uzbek State University of Physical Education and Sport, Tashkent, Uzbekistan; Research Institute of Epidemiology, Microbiology and Infectious Diseases, Tashkent, Uzbekistan

**Author notes:** **Corresponding author** Svetlana Osipova, MD, PhD, DSci, Address: Department of Immunology of Parasitic Diseases, Research Institute of Epidemiology, Microbiology and Infectious Diseases, Zakovat Str. 2, Uchtepa district, Tashkent city, Uzbekistan.

**Keywords:** vitamin D, elite athletes, water sport, cytokines, respiratory infection

## Abstract

**BACKGROUND:** The aim of this study is to identify seasonal changes in total 25(OH) vitamin D (VD) concentrations and determine its influence on acute upper respiratory tract infection (URTI) morbidity among elite athletes engaged in water sports.

**METHODS:** The study was planned as a prospective, non-interventional, observational study. Study participants included 40 elite athletes and 30 control individuals. Serum levels of 25(OH) VD and TNF-α, IFN-γ, IL-4 and IL-6 were detected by ELISA technique. Morbidity and frequency of acute URTI in participants were determined by self-reported questionnaire during the year.

**RESULTS:** The predominance of VD insufficiency was found in both groups of elite athletes and in the control individuals. Prevalence of VD insufficiency/deficiency depends on the season, but independently on the season the highest values were observed among athletes. VD sufficiency was detected in 30% and 13.3% of the control individuals in August and February and only in 10% of swimmers in August. More than 3 episodes of URTI were detected only in the elite athletes in winter-spring. The elevated level of TNF-α, IL-4, IL-6 was detected in all athletes, but more expressed increase was observed in swimmers.

**CONCLUSIONS:** VD insufficiency is quite pronounced among elite athletes engaged in synchronized swimming and in swimmers. It is accompanied with a decrease of IFN-γ, increase of TNF-α, IL-4 and IL-6 level, and elevation of URTI morbidity. Seasonal monitoring and correction of the VD level for normalization of cytokine profile and decrease of URTI morbidity is definitely advised.

## Introduction

The previous decade has been marked with the discovery of many new physiological functions of vitamin D (VD) as well as the documentation of epidemic VD deficiency and insufficiency worldwide.^1–4^ VD does not only influence the musculoskeletal health and mineral homeostasis but it also affects cardiovascular, endocrine, nervous, immune and mental functions,^5^ thus it is of considerable importance for the general population and especially for both physically active people and elite athletes in achievement of a physical peak condition and a high performance.^6^ VD status in athletes has been found to be variable and depending on outdoor training time, skin color, diet features, age and geographic location.^7–10^

VD concentration associates with incidences of upper respiratory tract infections (URTI): greater proportion of athletes maintaining circulating concentration of the blood serum 25(OH) VD <30 nmol/l presented more episodes of URTI as well as severity of symptoms than those with 25(OH) VD concentration >120 nmol/l.^11–13^ VD upregulates gene expression of antimicrobial peptides, which are important regulators in innate immunity and also downregulates expression of inflammatory cytokines such as TNF-α and IL-6,^14–15^ converting monocytes to macrophages, which in turn produce more inflammatory cytokines. Evidence was presented that influenza epidemics and other wintertime infections may be related to seasonal deficiencies of antimicrobial peptides secondary to VD deficiency.^16^ Despite a lot of studies of influence VD supplementation on incidence of URTI and flu, the results are contradictory.^17–20^

Water sports are characterized by certain specific features. Synchronized swimming combines speed, power and endurance. Athletes must train and compete spending a great amount of time underwater.^21^ URTI is one of the most common diseases in humans, with a high prevalence in athletes,^22–24^ and athletes engaged in water sports are of a special risk. An insufficient period of recovery between successive training sessions and overtraining syndrome may cause the suppression of the immune system, especially respiratory immune functions^25–26^ and may lead to an increase of the illness risk, notably, for the greater incidence of URTI. It has also been demonstrated in adult athletes that strenuous training might have an increased risk of infection because of transient immune suppression.^27–28^ Chronic low dose exposure of chlorine can induce airway hyperresponsiveness and aggravate allergic Th2 inflammation.^29^ Exposure to these chlorine-based chemical compounds can be manifested as clinical symptoms of upper respiratory dysfunction (chronic rhinitis, sneezing, irritated nasal sinuses, runny nose, etc.).^30^ It could contribute to increased susceptibility to URTI. Besides, inadequate nutrition status during heavy exercise can impair the respiratory immune functions.^31^ A well-designed training program with an adequate nutritional supplementation is needed not only to prevent the respiratory infections, but also to optimize the performance.^32–33^

The study of VD level and its impact on the frequency and susceptibility to URTI among elite athletes engaged in water sports is of great interest. Such studies have not been carried out in Central Asia countries previously as well as the prevalence of VD deficiency in the population.

The aim of the present study is to identify seasonal changes in total 25(OH) VD concentrations and determine its influence on acute URTI morbidity among elite athletes engaged in water sports.

## Materials and methods

### Study center

The prospective, non-interventional, observational study was conducted on the basis of the Uzbek State University of Physical Education and Sport and Research Institute of Epidemiology, Microbiology and Infectious Diseases, Tashkent, Uzbekistan, during the period from January 2017 till May 2018.

### Ethics approval

The study was approved by the Medical Ethics Committee of the Ministry of Health of the Republic of Uzbekistan in accordance with the Declaration of Helsinki. All participants provided written, informed consent after being informed about the protocol and purpose of the study. The trial is registered at the US National Institutes of Health (ClinicalTrials.gov) #NCT03623763.

### Study participants

Study participants included 20 elite athletes engaged in synchronized swimmers and 20 athletes engaged in swimming (all females) at the age of 19-24 years. For comparison of frequency of acute URTI, VD and cytokines levels the control group included 30 healthy individuals of the same sex and age without manifestations of diseases. All the participants were residents of Uzbekistan. Participants were required medical examination and to complete a comprehensive health screening questionnaire prior to the start of the study.^34^ Elite athletes could be included if they were currently healthy (with no health problems or infection symptoms within the previous two weeks) and engaged in regular sports training. Participants representing one or more of the following criteria were excluded from participation: smoking or use of any medication, suffering from or had a history of chronic infectious and noninfectious diseases (cardiac, hepatic, renal, pulmonary, neurological, gastrointestinal, hematological or psychiatric).

Frequency, duration and severity of acute URTI episodes and the data on the features of the diet during the year in elite athletes and the control individuals were determined by self-reported questionnaire, which was completed in February and August. The study was provided in February and August taking into consideration continental climate features of Uzbekistan (hot summer and relatively cold winter). On evidence derived from the Centre of Hydrometeorological Service of the Republic of Uzbekistan summer temperatures often surpass 40° C; winter temperatures average about 2° C, but may fall rarely as low as −40° C. The hottest period continues from June 25 until August 5. The coldest period continues from the 26 of December to 5 of February. The number of sunny days in a year is more than 300.

Criteria of acute URTI severity are shown in table 1.

**Table 1.**
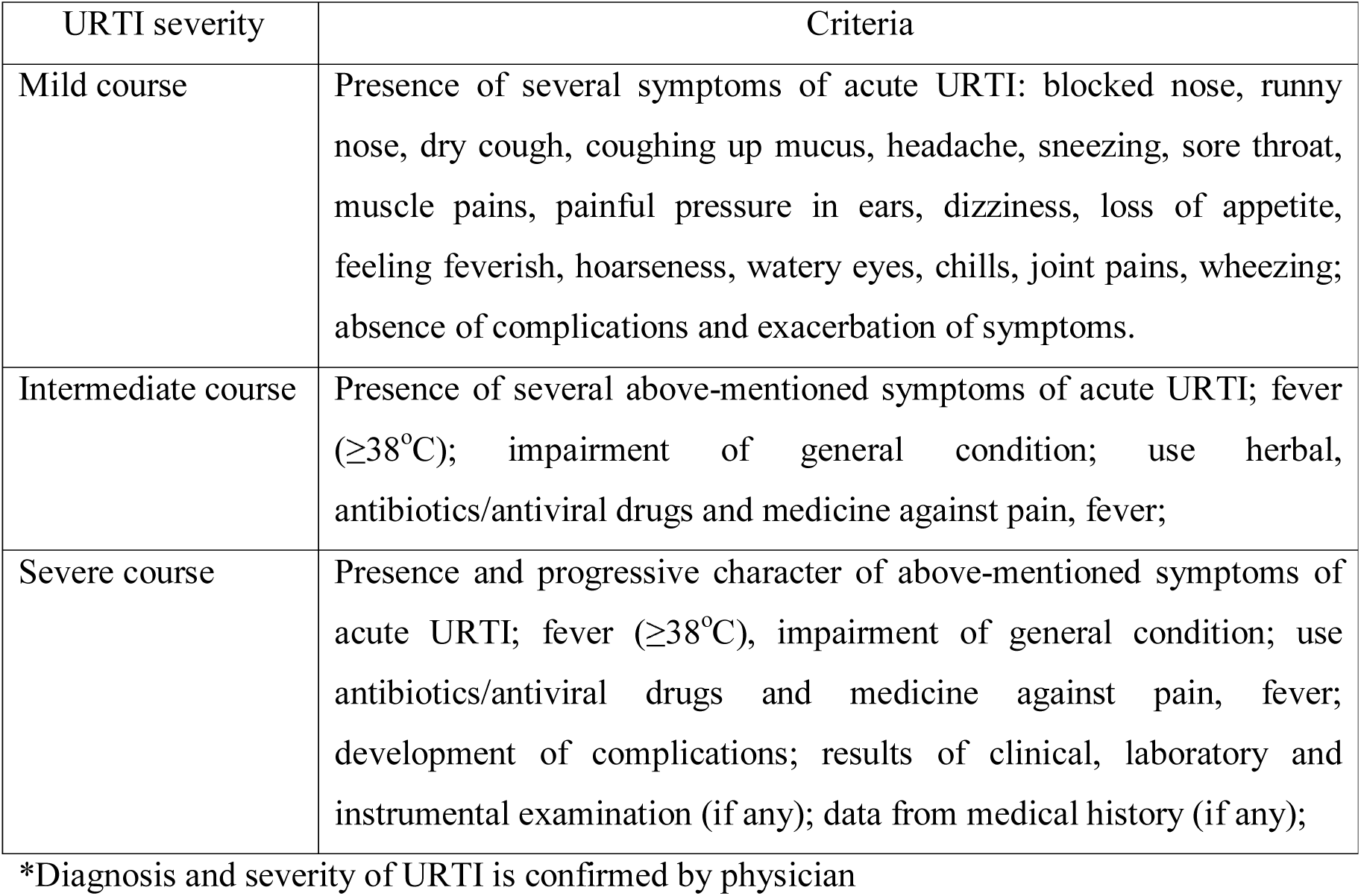
Criteria for diagnosis and assessing the severity of acute URTI*

| URTI severity | Criteria |
| --- | --- |
| Mild course | Presence of several symptoms of acute URTI: blocked nose, runny nose, dry cough, coughing up mucus, headache, sneezing, sore throat, muscle pains, painful pressure in ears, dizziness, loss of appetite, feeling feverish, hoarseness, watery eyes, chills, joint pains, wheezing; absence of complications and exacerbation of symptoms. |
| Intermediate course | Presence of several above-mentioned symptoms of acute URTI; fever ($\geq 38^{\circ}\text{C}$); impairment of general condition; use herbal, antibiotics/antiviral drugs and medicine against pain, fever; |
| Severe course | Presence and progressive character of above-mentioned symptoms of acute URTI; fever ($\geq 38^{\circ}\text{C}$), impairment of general condition; use antibiotics/antiviral drugs and medicine against pain, fever; development of complications; results of clinical, laboratory and instrumental examination (if any); data from medical history (if any); |

\*Diagnosis and severity of URTI is confirmed by physician

### Immunological tests

Participants were required to abstain from any strenuous physical activity for 24 h and fasting for at least six hours before coming to the laboratory. Five milliliters of peripheral venous blood was taken (after 8-12 hours of fasting) from each participant and were collected into Human Tube Serum Gel – C/A for ELISA. All blood samples were collected in February and August. Serum levels of 25(OH) VD (DIAsource kit, Belgium and LLC kit) and TNF-α, IFN-γ, IL-4 and IL-6 were detected by ELISA technique (Vector-Best, Novosibirsk, Russia) according to the manufacturer’s protocol. The performers of immunological tests did not have access to any information about an individual under examination. All information was blinded.

### Classification of the VD level

The serum VD level was classified as reported by Holick (2007).^4^ Levels of VD ≤20, 21–29, ≥30–150, and >150 ng/ml were considered as VD deficiency, VD insufficiency, VD sufficiency and VD intoxication, respectively.

### Statistical analysis

Data analysis was performed with the program Origin 6.1 (OriginLab, Northampton, MA). Results are expressed as mean±standard error (SEM) for continuous variables and number (percentage) for categorical data. For numerical variables the independent/paired t test was used. The P value <0.05 was considered as statistically significant.

## Results

40 elite athletes and 30 control individuals were enrolled in the study. The elite athletes were divided into two groups: swimmers (n=20) and synchronized swimmers (n=20). Forty-two were excluded as 19 participants refused to participate, and 23 did not meet the inclusion criteria (Figure 1). Individuals of the control group were matched by the baseline characteristics to the elite athletes. Analysis of baseline characteristics didn’t show any significant differences between elite athletes and the control individuals (Table 2).

**Figure 1.** The selection process of participants.

**Table 2.** Baseline characteristics of participants

| Characteristics | Swimmers<br>(n=20) | Synchronized<br>swimmers<br>(n=20) | Control<br>individuals<br>(n=30) |
| --- | --- | --- | --- |
| Age in years (SEM) | 20.3 (0.6) | 21.1 (1.2) | 19.8 (1.7) |
| Stature in m (SEM) | 1.674 (0.03) | 1.704 (0.02) | 1.654 (0.06) |
| Body mass in kg (SEM) | 59.6 (5.3) | 62.1 (3.1) | 51.8 (5.1) |
| BMI in kg/m<sup>2</sup> (SEM) | 21.2 (3.7) | 21.4 (2.1) | 18.9 (2.7) |
| Skin phototypes-no.(%±SE) |  |  |  |
| Fair skin, blue eyes | 7 (35±10.6) | 6 (30±10.2) | 12 (40±8.9) |
| Darker white skin | 8 (40±10.9) | 5 (25±9.6) | 6 (20±7.3) |
| Light brown skin | 4 (20±8.9) | 5 (25±9.6) | 4 (13.3±6.2) |
| Brown skin | 1 (5±4.8) | 4 (20±8.9) | 8 (26.7±8.0) |
| Years swimming, (SEM) | 5.7 (1.6) | 5.3 (0.9) | - |
| Amount of swimming, hr/week | 15 | 18 | - |
| Amount of gym training, hr/week | 6 | 6 | - |

During the analyzing of self-reported questionnaire diet features of elite athletes engaged in water sports and the control individuals were determined. The main daily using food products by athletes as well as the control individuals are milk, cream, cheese, rice, buckwheat, vegetable oil, butter, beef, lamb meat, chicken, fruit, vegetables, eggs, bread. Analysis of diet features of athletes revealed that 85% of athletes daily use food products with low content of VD, only 15% of athletes use food products with medium content of VD daily. No significant difference between the elite athletes and the control individuals in diet features was found.

Figure 2 A illustrates the predominance of VD insufficiency in both groups of athletes regardless of the season, especially in synchronized swimmers (100%) in comparison with the control individuals (63.3%) (P≤0.05). It should be noted that athletes engaged in synchronized swimming finished 30 days training in the outdoor swimming pool one week before blood serum test samples collection in August. However VD sufficiency was not found in any case. Training of swimmers took place in indoor swimming pools. VD sufficiency was determined in 10% of swimmers; VD deficiency was estimated in the same number of swimmers. VD sufficiency was found in 30% of the control individuals, but 6.6% of them were VD deficient. Figure 1 B demonstrates that in February an expressed decrease of VD level (deficiency) was detected in 30% and 20% of athletes engaged in swimming and synchronized swimming, respectively. The number of individuals with VD sufficiency in the control group was higher than in athletes.

**Figure 2.**
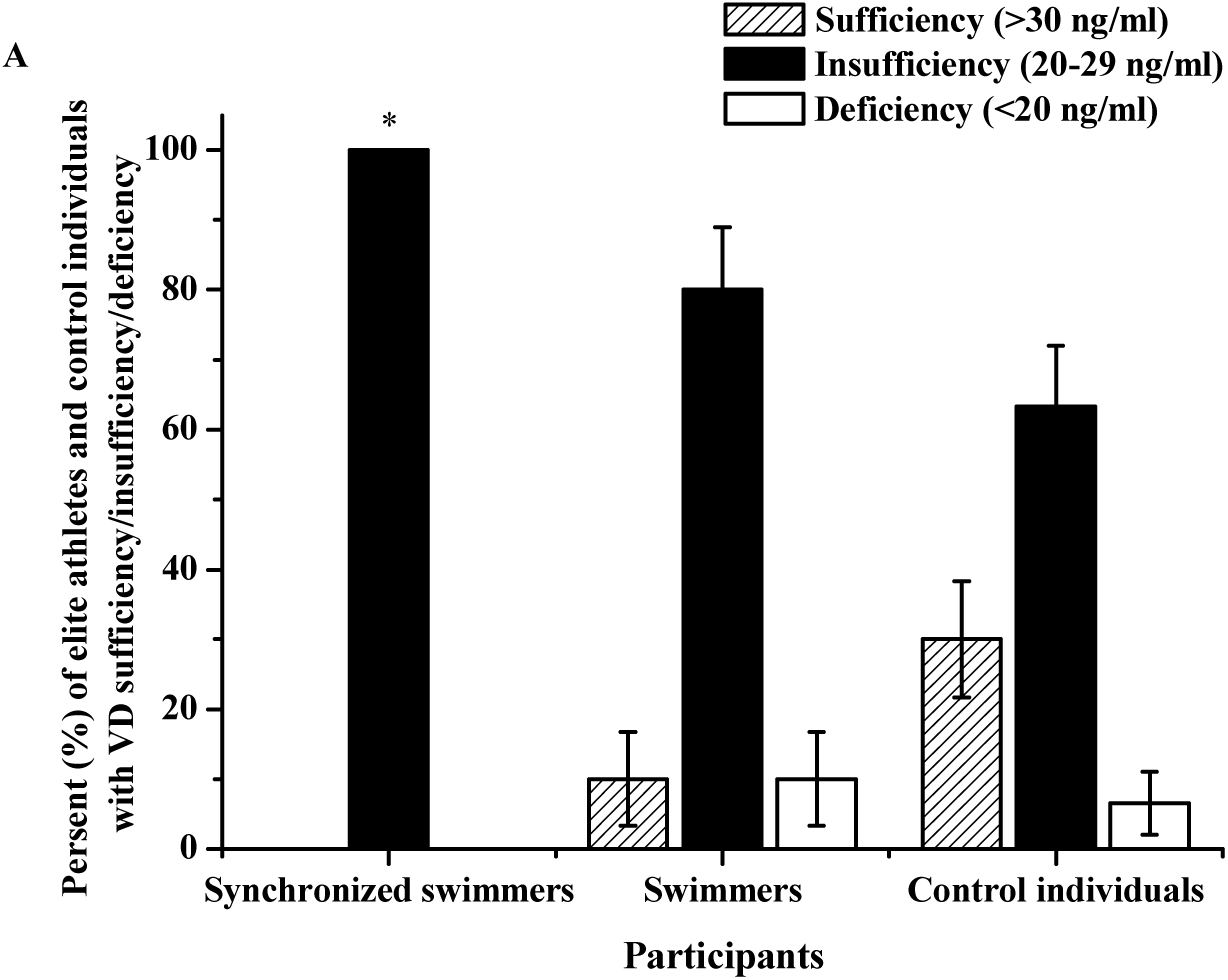
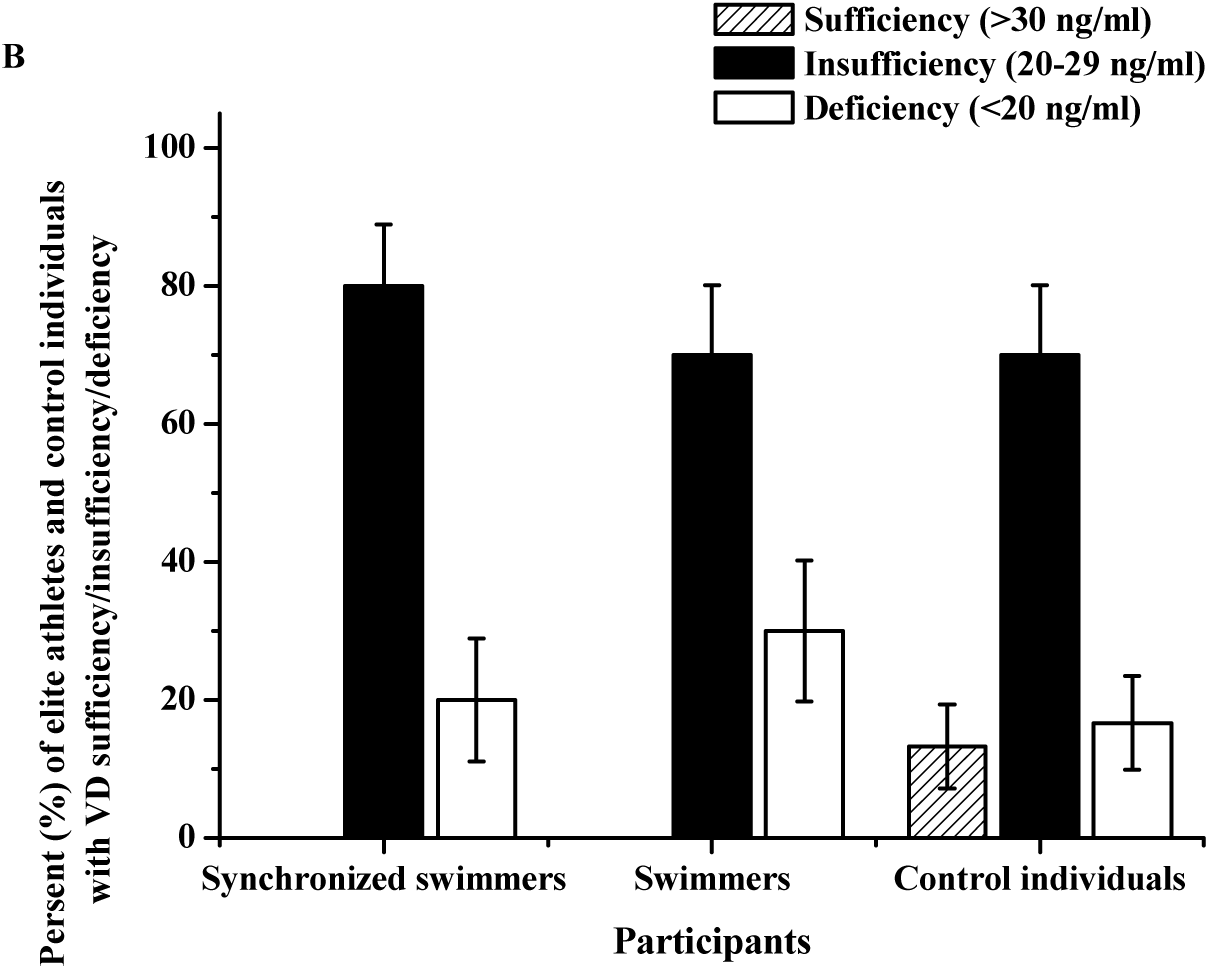
The level of serum 25(OH) VD in participants engaged in synchronized swimming, swimming and the control individuals in August (A) and February (B). A) Sufficiency of serum 25(OH) VD was detected in 10±6.7% of swimmers and 30±8.3% of control individuals; insufficiency of serum 25(OH) VD was detected in 100% of synchronized swimmers, 80±8.9% of swimmers and 63.3±8.7% of the control individuals; deficiency of serum 25(OH) VD was detected in 10±6.7% of swimmers and 6.6±4.5% of the control individuals; B) Sufficiency of serum 25(OH) VD was detected only in 13.3±6.1% of the control individuals; insufficiency of serum 25(OH) VD was found in 80±8.9% of synchronized swimmers, 70±10.2% of swimmers and 70.0±8.3% of the control individuals; deficiency of serum 25(OH) VD was detected in 20±8.9% of synchronized swimmers, 30±10.2% of swimmers and 16.7±6.8% of the control individuals; *significant difference with the control individuals (P<0.05)

The highest morbidity with acute URTI has been found in swimmers (55%) and synchronized swimmers (50%) during post-competition period. During the training process and pre-competition periods morbidity with acute URTI of swimmers (35% and 30%) and synchronized swimmers (30% and 20%) was not significantly differed. Distribution of elite athletes by according to duration of acute URTI shows the higher percentage of elite athletes with duration of URTI within 7-14 days and up to 14 days in post-competition period in comparison with training process and pre-competition periods. Percentage of elite athletes with acute URTI duration less than 7 days was equal in different periods (training process, pre-competition and post-competition periods). Severe course of URTI has been found only in swimmers (5%) in post-competition period. High frequency of intermediate course of URTI has been found in swimmers (30%) and synchronized swimmers (30%) in post-competition period. During the training process period intermediate course of URTI has been found in 15% of swimmers and 20% of synchronized swimmers. Only in 10% of swimmers intermediate course of URTI was observed. Apart of periods of training process and competition the percentage of elite athletes with mild course of URTI during the year was varied within 10-20% (Table 3).

**Table 3.**
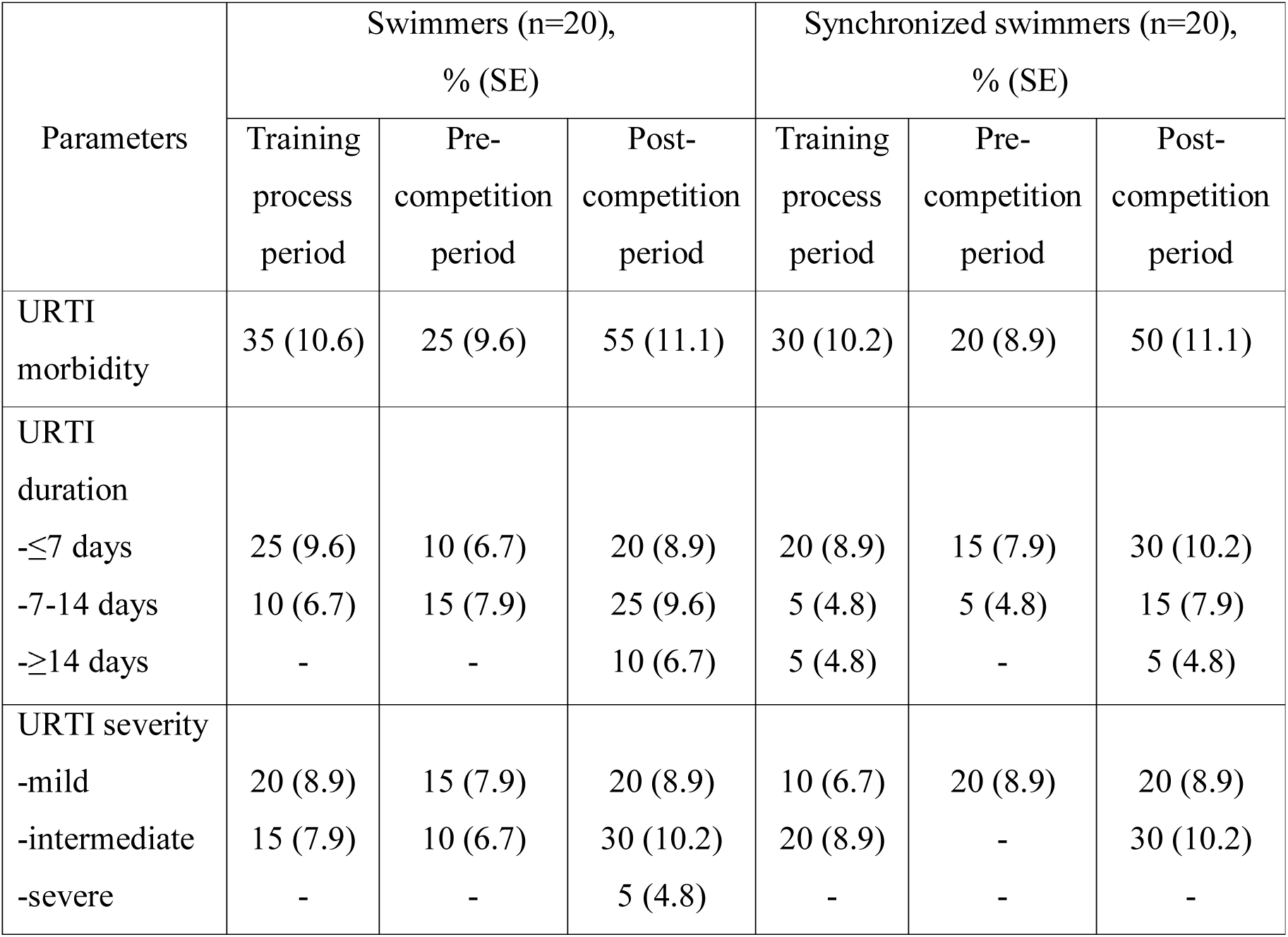
Morbidity, duration and severity of acute URTI during the training process, pre- and post-competition period among water sports elite athletes

Table 4 shows the frequency of URTI during summer-autumn and winter-spring periods among athletes and the control individuals. In both groups of athletes and in the control group individuals URTI were observed more frequently in winter-spring period. However, more than 3 episodes of acute URTI were detected only in elite athletes in winter-spring. 2-3 episodes of URTI regardless of the season were significantly more frequently detected in both groups of elite athletes in comparison with the control individuals (P≤0.05). Absence of URTI was observed in a very low percentage of elite athletes versus the control individuals.

**Table 4.** The frequency of URTI during summer-autumn (sum-aut) and winter-spring (win-spr) periods among athletes and the control individuals

| The number<br>of episodes<br>of URTI | The frequency of URTI among participants during summer-autumn and<br>winter-spring periods, % (SE) |  |  |  |  |  |
| --- | --- | --- | --- | --- | --- | --- |
|  | Synchronized<br>swimmers<br>(n=20) |  | Swimmers<br>(n=20) |  | Control individuals<br>(n=30) |  |
|  | sum-aut | win-spr | sum-aut | win-spr | sum-aut | win-spr |
| Absence of URTI | 40 (10.9)* | - | 25 (9.6)* | 5 (4.8)* | 80 (7.3) | 36.7 (8.7) |
| ≤2 episodes | 35 (10.6)* | 30 (10.2) | 15 (7.9) | 25 (9.6)* | 16.7 (6.8) | 40 (8.9) |
| 2-3 episodes | 25 (9.6)* | 45 (11.1)* | 60 (10.9)* | 40 (10.9)* | 3.3 (3.2) | 23.3 (7.7) |
| 3-4 episodes | - | 25 (9.6) | - | 20 (8.9) | - | - |
| ≥5 episodes | - | - | - | 10 (6.7) | - | - |

\* - significant difference with the control individuals (P<0.05)

Results of cytokines detection performed in August and February are presented in Table 5. Level of TNF-α was significantly elevated in all athletes in comparison with the control individuals, but in athletes engaged in synchronized swimming it was lower, than in swimmers. The elevated level of IL-4 and IL-6 was detected in all athletes, but more expressed increase was observed in swimmers. A decrease of IFN-γ was observed in all athletes, however the lowest values significantly differed from the control values were found in swimmers.

**Table 5.** The level of serum cytokines (TNF-α, IFN-γ) in elite athletes

| Cohort<br>under study | Cytokines (pg/ml), (SEM) |  |  |  |  |  |  |  |
| --- | --- | --- | --- | --- | --- | --- | --- | --- |
| | TNF-α | | IFN-γ | | IL-4 | | IL-6 | |
|  | Aug. | Feb. | Aug. | Feb. | Aug. | Feb. | Aug. | Feb. |
| Synchronized swimmers (n=20) | 12.0 (2.1)* | 17.9 (1.9)* | 15.6 (1.9) | 12.7 (1.3) | 7.5 (1.4)* | 5.5 (1.2) | 28.1 (4.6) | 35.9 (5.8) |
| Swimmers (n=20) | 16.1 (2.6)* | 20.4 (3.7)* | 10.0 (1.2) | 9.1 (0.8) | 9.0 (2.11)* | 12.9 (3.21)* | 31.0 (6.1) | 48.9 (8.1)* |
| Control individuals (n=30) | 4.4 (1.7) | 5.1 (1.2) | 19.4 (2.2) | 17.5 (1.8) | 1.7 (0.9) | 2.7 (0.9) | 24.1 (1.0) | 20.3 (0.8) |

\* - significant difference with the control individuals (P<0.05)

## Discussion

The study is the first experience of detection of VD level and its influence on acute URTI morbidity among elite athletes engaged in water sports in Uzbekistan. The predominance of VD insufficiency was found in both groups of elite athletes and the control individuals despite the large number of sunny days in the country (>300 days per year). Diet features of study participants are associated with regional diet habits and geographical location. Access to sea foods containing the high content of VD is almost absent in Uzbekistan. Other food products containing the high content of VD also rarely using in daily food ration. According to analysis of self-reported questionnaires insufficient dietary VD intake has been found in elite athletes. Higher prevalence of VD insufficiency/deficiency in both groups of athletes could be explained by the influence of significant physical load, prolonged time in indoor trainings and diet features. It is important to note that the current study population consisted mainly of elite athletes with a white skin colour and that the seasonal variation of VD might be different in athletes with a darker skin colour. Our data confirm literature data about the high prevalence of inadequate VD level among athletes and the general population.^35–38^

The regulatory effect of 25(HO) VD on the immune system is well known. Active 25(HO) VD induces the synthesis of antimicrobial peptides by natural killers and respiratory tract epithelial cells,^39^ as well as upregulated S100 protein and calprotectin levels were identified under the effect of active 25(HO) VD.^40^ In the case of a VD deficiency, the immune response is disrupted and leukocyte chemotaxis is reduced.^41^ The rate of infection diseases rises due to disrupted immunity. Yim and colleagues (2007) revealed that antimicrobial protein cathelicidin synthesis was impaired in the bronchial epithelial cells in patients with frequent respiratory tract infection.^42^ Deterioration of synthesis of the 25(HO) VD dependent immune regulator proteins such as cathelicidin, defending, S100, and calprotectin may clarify the increased URTI in patients with 25(HO) VD deficiency. Nasal mucociliary clearance time was significantly increased in patients with VD deficiency which might be one of the pathophysiologic pathways at increased URTI.^43^ Some studies propose that 25(HO) VD may be effective in protecting against respiratory tract infections.^44,45^ In spite of the literature, the pathophysiologic pathway of the increased rate of URTI still remains unclear. The most known hypothesis is an impaired immune response in patients with VD deficiency. However, this is not well documented and not enough to explain pathophysiologic pathways.

URTI are thought to impair athletic performance; with athletes and coaches placing a high priority on preventing illness.^46^ It has been demonstrated that the incidence of URTI was associated with reduced training load by 24% in endurance sports athletes, and this insufficient training load may increase the injury rates and decrease the physical capacity.^47,48^ We found the higher TNF-α level in swimmers, which could be explained by heavier training load. The elevated level of IL-4 and IL-6 was detected in all athletes, but more expressed increase was observed in swimmers. Kapilevich et al. (2017) showed that the level of circulating IL-6 is connected with character of physical load (dynamic or static).^49^ The higher elevation of IL-6 level in swimmers can be connected with significant volume and intensity of training. The level of IL-6 is known to be the first response to the volume and type of exercise.^50^ Increased IL-4 and IL-6 level consequently contributes to progression of URTI.^51^

A decrease of IFN-γ was observed in all athletes and the lowest values were found in swimmers. Little is known about the connection of IFN-γ level and VD insufficiency in athletes. It was shown that minimal IFN-γ values were detected together with a VD deficiency.^52^ In athletes with a VD sufficiency concentration of circulating IL-10 and IFN-γ are significantly higher than in athletes with a VD deficiency.^51^ Identified trend to elevation of URTI morbidity in athletes with VD deficiency and lower values of VD insufficiency, changes of the cytokine profile of our results are in accordance with the data by Greiller & Martineau (2015) on the role of VD in resistance to URTI.^53^ In total, our data are in accordance with information on the possible connections of VD deficiency/insufficiency with features of cytokine profile and URTI morbidity. Further trials are needed to determine the most appropriate VD supplementation regime in athletes engaged in water sports in general and in particularly for Uzbekistan.

Our study has several limitations. Firstly, the number of participants who adequately completed the self-reported questionnaires and give consent to participation was lower than anticipated. Another limitation is that the diagnosis of acute URTI was only symptomatically and verification of the etiology of diseases was not done due to the difficulty of diagnosis of wide spectrum of infectious agents of acute URTI and non appearance of participants in pre- and post-competition periods and mild course of disease.

## Conclusion

VD insufficiency is quite pronounced in elite athletes engaged in synchronized swimming and in swimmers. It is accompanied with decrease of IFN-γ, increase of TNF-α, IL-4 and IL-6 level, and elevation of URTI morbidity. Seasonal monitoring and correction of VD level for normalization of cytokine profile and decrease of URTI morbidity is definitely advised.

## Author contributions

Conceptualization: SO, FK, JU; Methodology: SO, AT, ND; Software: AT, ND, JU; Validation: SO, FK; Formal analysis: SO, AT, FK; Investigation: AT, JU; Data curation: SO, FK; Writing (original draft preparation, review and editing): SO, AT, JU; Visualization: AT, ND, JU; Supervision: SO, FK; All authors participated in the conception and design of the study and the interpretation of data, and all approve the final manuscript for publication.

